# Biological classification with RNA-Seq data: Can alternative splicing enhance machine learning classifier?

**DOI:** 10.1101/146340

**Authors:** Nathan T. Johnson, Andi Dhroso, Katelyn J. Hughes, Dmitry Korkin

## Abstract

The extent to which the genes are expressed in the cell can be simplistically defined as a function of one or more factors of the environment, lifestyle, and genetics. RNA sequencing (RNA-Seq) is becoming a prevalent approach to quantify gene expression, and is expected to gain better insights to a number of biological and biomedical questions, compared to the DNA microarrays. Most importantly, RNA-Seq allows to quantify expression at the gene and alternative splicing isoform levels. However, leveraging the RNA-Seq data requires development of new data mining and analytics methods. Supervised machine learning methods are commonly used approaches for biological data analysis, and have recently gained attention for their applications to the RNA-Seq data.

In this work, we assess the utility of supervised learning methods trained on RNA-Seq data for a diverse range of biological classification tasks. We hypothesize that the isoform-level expression data is more informative for biological classification tasks than the gene-level expression data. Our large-scale assessment is done through utilizing multiple datasets, organisms, lab groups, and RNA-Seq analysis pipelines. Overall, we performed and assessed 61 biological classification problems that leverage three independent RNA-Seq datasets and include over 2,000 samples that come from multiple organisms, lab groups, and RNA-Seq analyses. These 61 problems include predictions of the tissue type, sex, or age of the sample, healthy or cancerous phenotypes and, the pathological tumor stage for the samples from the cancerous tissue. For each classification problem, the performance of three normalization techniques and six machine learning classifiers was explored. We find that for every single classification problem, the isoform-based classifiers outperform or are comparable with gene expression based methods. The top-performing supervised learning techniques reached a near perfect classification accuracy, demonstrating the utility of supervised learning for RNA-Seq based data analysis.

## Introduction

Ever since the intrinsic role of RNA was proposed by Crick in his Central Dogma [1], there has been a desire to accurately annotate and quantify the amount of RNA material in the cell. A decade ago, with the introduction of RNA sequencing (RNA-Seq) [2], it became possible to quantify the RNA levels on the whole genome scale using a probe-free approach, gaining insights into cellular and disease processes and illuminating the details of many critical molecular events such as alternative splicing, gene fusion, single nucleotide variation, and differential gene expression [3]. The basic assessment of RNA-Seq is focused on utilizing the data for differential gene expression between the groups of biological importance [4]. However, there are additional patterns that can be elucidated from the same raw sequencing data by extracting the expression levels of the alternatively spliced isoforms [5].

Alternative splicing (AS) of pre-mRNA provides an important means of genetic control [6, 7]. It is abundant across all eukaryotes and even occurs in some bacteria and archaea [8-10]. AS is defined by the rearrangement of exons, introns, and/or untranslated regions that yields multiple transcripts [11]. Furthermore, 86-95% of multi-exon human genes is estimated to undergo alternative splicing [12]. Genes tend to express many isoforms simultaneously, 70% of which encode important functional or structural changes for the protein [12]. RNA-Seq data encompasses expression at both gene and isoform levels: the gene-level expression amounts to the combined expression of all isoforms associated with the particular gene. It has been previously demonstrated that the gene-level expression is an excellent indicator of the tissue of origin as well as certain cancer types [13-17]. However, the isoform-level expression has been suggested to provide a more precise measurement of gene product dosage [5]. Differential AS depends on many factors, including the epigenetic state, genome sequence, RNA sequence specificity, activators and inhibitors from both, proteins and RNAs, as well as post-translational modification [6, 18-20]. These diverse mechanisms control AS to obtain developmental, cell-type, and tissue-specific expression. Furthermore, the patterns driven by AS and specific to cancer and other diseases have been recently identified [21, 22].

Machine learning tools developed over the last several decades have significantly advanced the analysis of the vast amount of next generation sequencing and microarray expression data by discovering the biologically relevant patterns [23-25]. Previous studies have utilized unsupervised and supervised machine learning techniques on the microarray gene expression data with variable success rates [26, 27]. Along with the individual approaches [28], large-scale comparative studies have been carried out [29, 30]. Some studies evaluated both basic and advanced clustering techniques, such as hierarchical clustering, k-means, CLICK, dynamical clustering, and self-organizing maps, to identify the groups of genes that share similar functions or genes that are expressed during the same time point of a mitotic cell cycle [17, 31, 32]. Other studies compared the ability to perform disease/healthy sample classification tasks by state-of-the-art supervised methods, such as Support Vector Machines (SVM), Artificial Neural Nets (ANN), Bayesian Networks, Decision Trees, and Random Forest classifiers [30].

When it comes to the biological classification, the RNA-Seq data present an attractive alternative to microarrays, since it is possible to quantify all RNA present in the sample without the need of the *a priori* knowledge. With RNA-Seq rapidly replacing microarrays, it is necessary to assess the potential of the supervised machine learning methodology applied to the RNA-Seq data across multiple datasets and biological questions [33]. Recently, there have been limited studies that have assessed RNA-Seq data with supervised and unsupervised machine learning techniques [34]. However, these studies utilized RNA-Seq data by leveraging only gene-level expression data rather than more detailed transcript-level, or isoform-level, data available for the alternative splicing isoforms [6]. Most recently, a study analyzed the utility of RNA-Seq isoform-level data for the disease/non-disease phenotype classification of the samples, showing the advantage of the isoform expression data for the disease phenotype prediction task [35]. However the question of whether or not the utility of isoform-level expression presents a general trend across all main biological and biomedical classification tasks remains open.

This work aims to systematically assess how well state-of-the-art supervised machine learning methods perform in various biological classification tasks when utilizing either gene-level or isoform-level expression data obtained from the RNA-Seq experiments. The assessment is done from three different perspectives: (i) by analyzing RNA-Seq data from two organisms (rat and human), (ii) by using the increasingly difficult datasets, and (iii) by considering different technical scenarios. The datasets were analyzed using six supervised machine learning techniques, three normalization methods, and two RNA-Seq analysis pipelines. Altogether, the performance on 61 major classification problems that include 2,196 individual classification tasks were compared. The use of multiple datasets allows us to determine if the success of a classification task is due to the discovery of distinct biological patterns by a machine learning algorithm, or if it is due to biologically unrelated patterns caused by differences in library preparation and/or the lab source. Finally, we assess whether using the information on alternatively spliced isoforms presented in the form of transcript expression data can provide the higher classification accuracy, compared to the gene expression data.

## Results

The goal of this work is to examine the capabilities of supervised machine learning methods in performing biological classification based on RNA-Seq data. Specifically, we analyzed whether the performance is influenced by (1) the power of the machine learning classifier, and/or (2) more detailed information extracted from the RNA-Seq data. In the first case, we assessed several supervised classifiers, ranging from the very basic methods to the state-of-the-art supervised classifiers, across three different normalization techniques. In the second case, we compared the same classifiers using either gene-level or isoform-level expression data. Together, the study setup utilized three RNA-Seq datasets, six supervised machine learning techniques, and three normalization protocols (Figure 1). Furthermore, each of the 61 classification problems was set to use the numerical features generated either from the gene-level or isoform-level expression data. To the best of our knowledge, this is the largest comparative analysis of biological classification tasks based on RNA-Seq data, performed to date.

**Figure 1.**
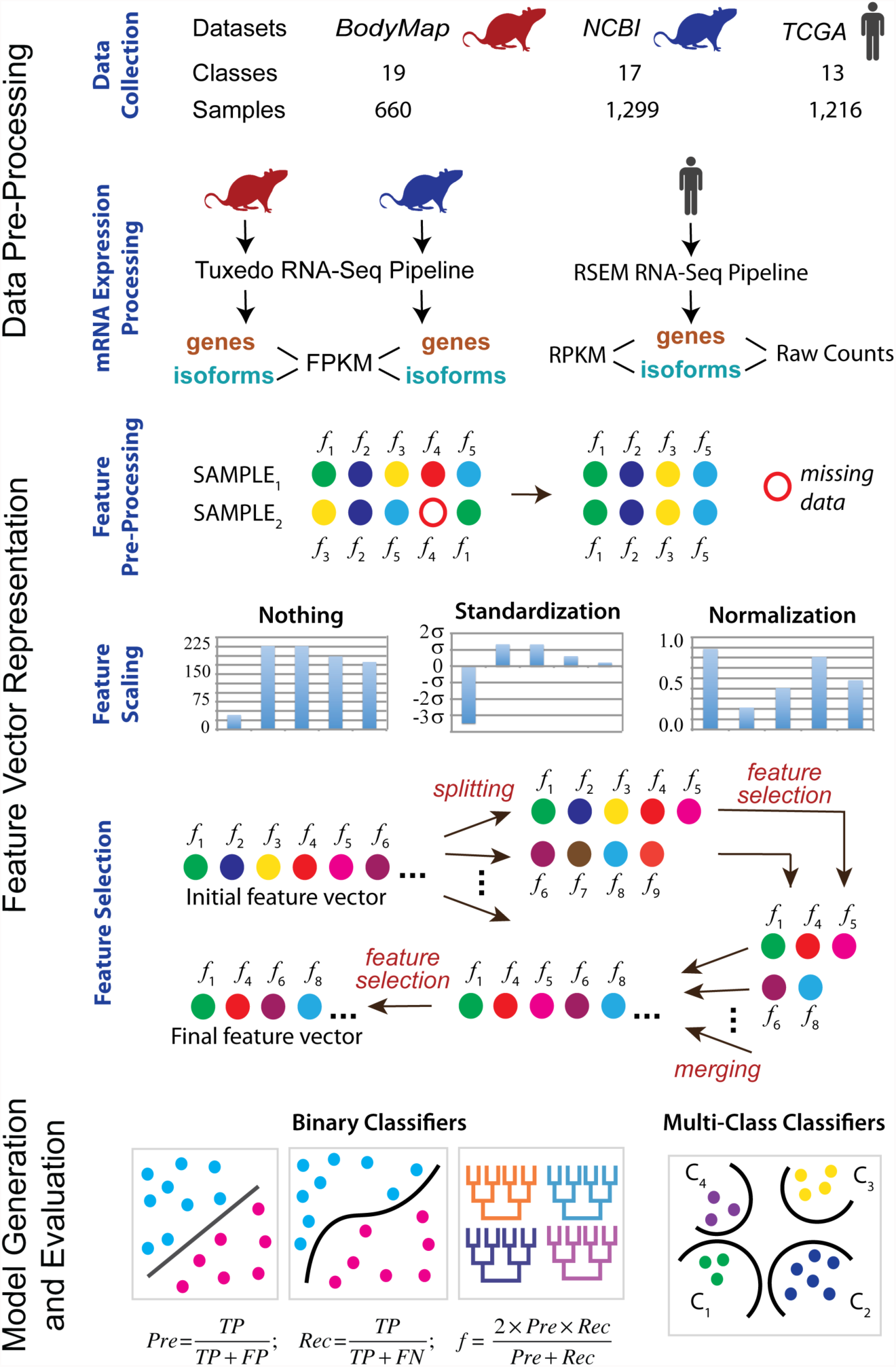
Overall computational pipeline used in this work. The samples from each of the three datasets are collected. The classification tasks are then defined. The expression data are processed for each sample at the gene and isoforms levels using two RNA processing pipelines and three different count measures. Next, feature pre-processing, scaling, and selection are done for each classification task. Finally, the binary as well as multiclass supervised classifiers are trained and tested.

### Classification Tasks Analyzed

Two categories of classification tasks were considered: normal phenotype and disease phenotype. In the first category, we determined whether it was possible to distinguish between age groups, sex, or tissue types in normal rats based on transcriptome analysis. The second category focused on classification tasks associated with breast cancer, with the main goal to differentiate between the pathological tumor stages. Both categories were analyzed using RNA-Seq data at the gene and isoform levels. Two types of classification were considered for each category of tasks: binary classification and multiclass, or multinomial, classification. These classification types center around two conceptually different classification problems. The binary classification distinguishes a sample as either belonging to the class or not. The multiclass classification distinguishes which class a specific sample belongs to. For example, for a binary tissue classification task, a sample can be classified as extracted from the brain tissue or not. In the context of a multiclass classification, the same sample is classified as extracted from exactly one of several tissue types.

### Dataset Statistics

Three datasets were used to carry out the classification tasks: two datasets for the normal phenotype classification tasks and one dataset for the disease phenotype classification tasks (Figure 1). The first dataset was obtained from the Rat Body Map, and is referred to as RBM dataset. It consisted of 660 normal rat samples whose transcriptomes were sequenced from the same rat strain and served a reference dataset for the community [36]. The transcriptomes were obtained at 40 M reads per sample on average. The data were evenly split between the male and female rats, four age groups, and eleven tissue types (Suppl. Figure S1). The four age groups included 2, 6, 21, and 101 weeks. The eleven tissue types included adrenal gland, brain, heart, thymus, lung, liver, kidney, uterus, testis, muscle, and spleen. All samples used the same library preparation protocol, sequencer, and were prepared by the same laboratory. As a result, the dataset was expected to have the least impact from the data inconsistency that arises from the non-biological sources, such as utilizing different sequencing protocols, instruments, and other parameters.

The second dataset, also used in the normal phenotypes classification tasks and referred to as NCBI dataset, included over 1,100 samples (Suppl. Figure S2) with the sequencing depth ranging between 6 and 116 M reads. This dataset was prepared by analyzing the collection of rat transcriptomes that were sequenced on Illumina Hi-Seq 2000 platform and were publically available from the NCBI SRA database [37]. The dataset was obtained from the sequencing experiments of 29 research projects. It contained highly variable transcriptomes due to the differences in library preparation, project goals, and rat strains (Suppl. Figure S10, Table S3). The classification tasks for the NCBI dataset were the same as for the RBM dataset, but with one modification. The age classification was modified from the four age groups into either embryo or adult age groups (Suppl. Table S1) and is described later in this section.

The last dataset included raw RNA-Seq data from 1,216 human breast cancer patients from the Cancer Genome Atlas (referred to as TCGA dataset) and was used in the disease phenotype classification tasks [38]. At the preprocessing stage, two RNA-Seq data normalization techniques were implemented and compared. Classification was performed to distinguish between the pathological cancer stages, as defined by the American Joint Committee on Cancer (AJCC) [39]. The AJCC breast cancer staging is based on size of tumors present within the breast, presence or absence of detection of metastases that are not within the breast, and the presence, size, and type of metastases within the lymph nodes. The patients were distributed roughly evenly across the four main cancer stages, but with high variability when considering sub-cancer stages (Suppl. Figure S11).

### Feature Selection and Analysis

The numerical features for this study represented either gene or isoform expression levels. As a result, the number of features ranged from 10,711 to 73,592, depending on the dataset and representation (Suppl. Table S1). Utilizing all features for a classification task greatly increases the computational complexity. Moreover, not all expressed genes or isoforms may be important for a given classification task; using the uninformative features during the training process could potentially decrease the accuracy of the classifier. To reduce the dimensionality of the feature space, a feature selection method [40] was applied in a classification-specific and dataset-specific manner, resulting in a significant reduction of features ranging from 107 to 735 folds (Figure 2A, Suppl. Figures S1-S8, Suppl. Tables S1-S2).

**Figure 2.**
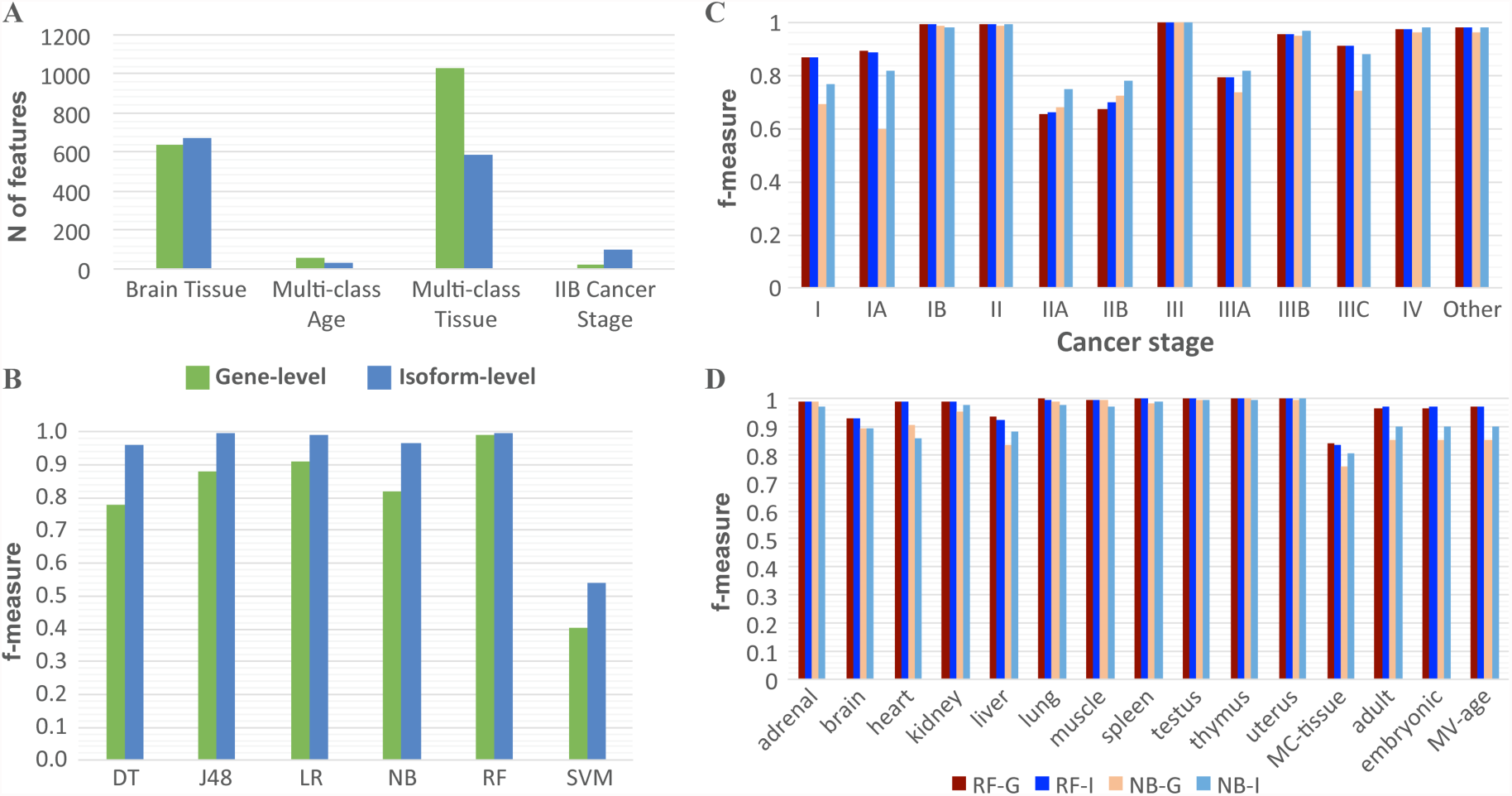
Overview of feature selection and the performance of classifiers using gene and isoform level expression data. **A**. Comparison of the number of features between gene and isoform after feature selection. Each classification task has the same number of features selected for each classifier at the gen-level and isoform-level. The four selected classes represent the four types of patterns seen between gene-level (green) and isoform-level classifiers. The Brain Tissue class is the most common pattern of feature selection. In general, more features are selected for isoform-level classifiers versus gene-level. **B.** Example of the variability of gene and isoform performance determined by f-measure across the six methods (DT = Decision Table, J48 = J48 Decision Tree, LR = Linear Regression, NB = Naïve Bayes, RF = Random Forest, SVM = Support Vector Machine). This example is from the RBM dataset for the Multi Age class without normalization. While there is a high degree of variability in performance, isoform-level classifiers consistently perform either comparably or better than gene-level classifiers. **C.** and **D.** Summary of the performance variability across classes for gene and isoform f-measure for the most frequent top and bottom performance methods (RF-G = Random Forest Gene, RF-I = Random Forest Isoform, NB-G = Naïve Bayes Gene, NB-I = Naïve Bayes Isoform). The data used in (C) is TCGA dataset and in (D) is NCBI dataset. MC = stands for multiclass.

Regardless of the classification task or dataset, the normalization of the RNA-Seq data did not make a significant difference on the choice of the selected features: variation in the numbers of selected features was less than 1% (Suppl. Figures S5-S8). An interesting observation, consistent across different tissue classification tasks, was that the number of features selected for the multiclass classification tasks was significantly greater than for a binary classification task. This observation should not be surprising, because the binary classification task is generally simpler than the multiclass classification task (Suppl. Table S2). However, in our case even if all features of the binary classification tasks related to a single multiclass classification task were combined, it would still not account for all features selected by the feature selection method for the multiclass task.

In many cases, the overall number of features selected for a binary classification task was the same or nearly the same, irrespective of whether the features were gene- or isoform-based (Suppl. Figures S1-S4). Does it mean that the features from the gene- or isoform-based approaches correspond to the same genes? Not always: the transcripts used for the selected features in a transcript-based, or isoform-based, classifier did not always originate from the genes that were selected for the corresponding gene-based classifier. Indeed, because 70% of isoforms were expected to encode different functional gene products [12], we expected cases where the gene expression features were not as specific as the corresponding isoform features. In general, there was a significant portion of 73,592 isoforms from 20,524 genes that corresponded to the same gene set (70-100%). However, there were several classification tasks, including multiclass tissue classification using NCBI dataset, where there was a lower percentage of such overlap (30%). Furthermore, there were several classification tasks, including multiclass age classification using RBM, multiclass tissue classification using GEO, and stage IIB classification using TCGA dataset, where the numbers of features that used either gene or isoform level of expression were significantly different, which was usually the case when a multiclass classification task was considered (Figure 1A, Suppl. Figures S1-S4, Suppl. Tables S1, S2). Another interesting observation was obtained when comparing the RBM and NCBI Rat datasets: the number of selected features was much smaller for the RBM dataset rather than for the NCBI Rat dataset (on average 231 versus 588), thus indicating the need for additional features to compensate for the increased data variability found within the NCBI Rat dataset.

### Overall Performance of Classifiers trained on Gene-based vs. Isoform-based data

Next, we hypothesized that because of the observed specificity of alternative splicing across tissues, ages, sexes, and between disease/normal phenotypes, training classifiers with the RNA-Seq data at the transcript, or isoform, level for the biological classification tasks could increase the classification accuracy [41]. Consistent with this hypothesis, the supervised learning classifiers that leverage the isoform-based data performed comparably or better than the classifiers trained on the gene data for all classification tasks (Figures 2B, 3). This observation also held true irrespective of the datasets used, normalization protocols, classification tasks, or supervised classifiers. The most frequently top performing methods were the random forest and logistic regression classifiers, whereas the worst performing method was typically naïve Bayes classifier (Figures 2B-D). However, the former approaches were not the most accurate ones for every single classification task, since in some cases naïve Bayes classifier was capable of outperforming all other methods tested (Stage IIA & IIB, Figure 2C). In general, the random forest classifier applied to the data without any normalization achieved 83-100% accuracy (Figures 2B-D).

**Figure 3.**
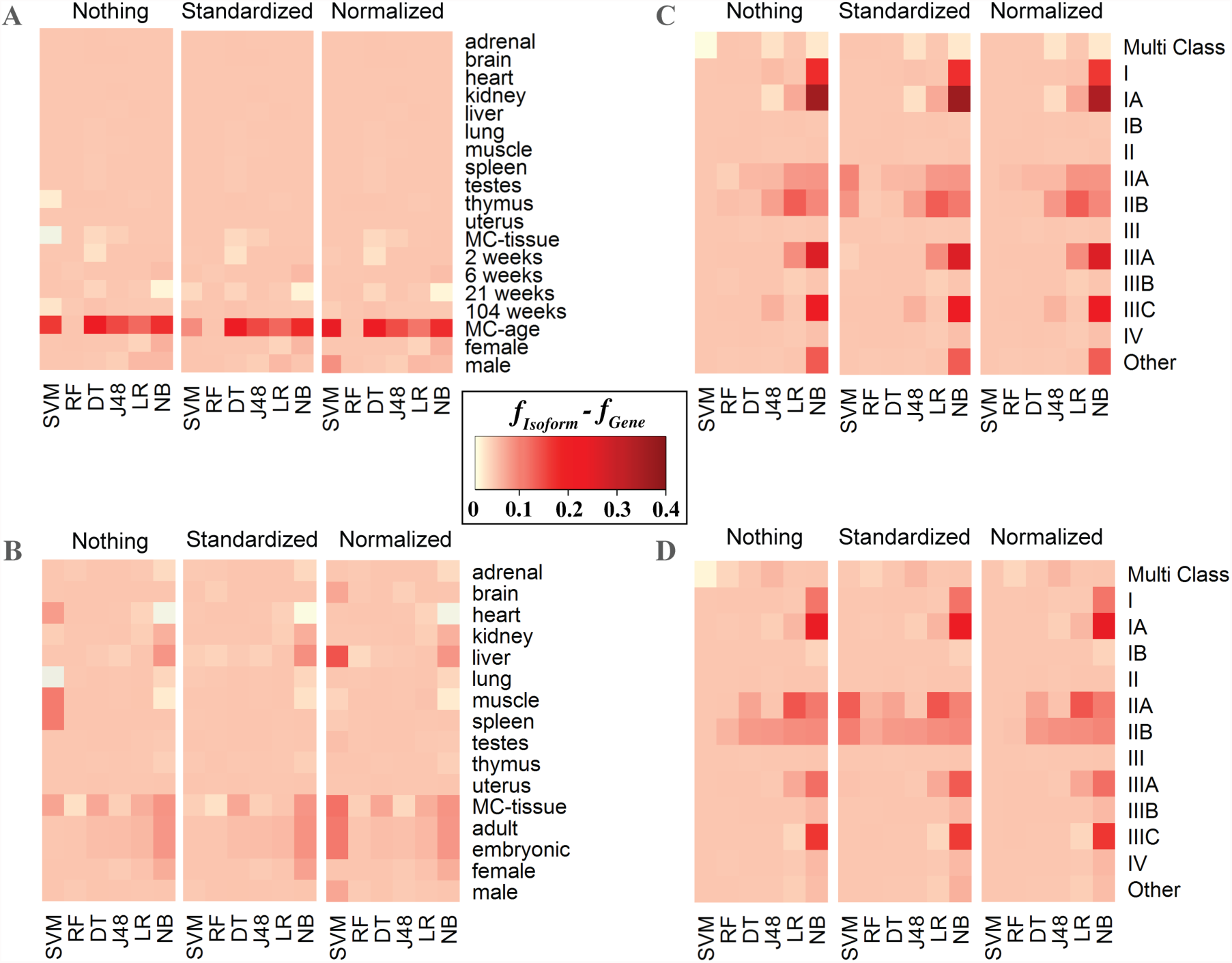
Heat map representation of the difference between Isoform and Gene *f*-measure across machine learning methods, classes, datasets, and normalization techniques. For the majority of classification tasks using isoform-level rather than gene-level expression data resulted in small to substantial increase of the performance accuracy, represented by *f*-measure values here. The bottom x-axis represents the machine learning techniques (DT = Decision Table, J48 = J48 Decision Tree, LR = Linear Regression, NB = Naïve Bayes, RF = Random Forest, SVM = Support Vector Machine). The y-axis represents the classes where MC stands for multiclass. The top x-axis represents normalization techniques including Nothing (no normalization), Standardized, and Normalized. Datasets for each panel are (A) RBM, (B) NCBI, (C) TCGA – log_2_ normalized counts, (D) TCGA – raw counts.

It was also observed that for 63% of classification tasks, the gene- and isoform-based methods performed with the same accuracy. For 37% of the classification tasks, the isoform-based methods performed better than the gene-based (more than 0.2 gain in f-measure value). The difference between the isoform-based and gene-based classification accuracies was particularly profound when comparing the classification results of naïve Bayes, which was one of the less accurate methods analyzed, while being among the fastest classifiers. However, we did not observe such a drastic difference, and sometimes no difference at all, when considering one of the most accurate classifiers, random forest, across all classification tasks. For instance, when comparing gene- and isoform-based classifiers for stage IA cancer using the raw count expression values and not performing any normalization protocols, the accuracy and f-measure values for naïve Bayes classifier ranged between 49.5%-76.4% and between 0.60-0.82, respectively, while for random forest the ranges were nearly identical (Figures 2C, D).

Regardless of using gene- or isoform-based approaches on different datasets, classification tasks, or supervised learning methods, it was observed that the three primary normalization techniques used when processing RNA-Seq data did not improve or reduce the accuracy. The only exception was the performance of SVM classifier employed by both, the isoform-based and gene-based, approaches: differences in the accuracy values between the various normalization techniques for some classification tasks were as high as 40.3 and 30.7%, respectively (Figure 5).

Another potential source of variability in the classifier performance was the difference in the protocols used by different studies. To determine whether the difficulty of classification task increased when using datasets from multiple laboratories rather than from a single one, the classification accuracies between the two rat datasets were compared for each binary or multiclass classification task. Not surprisingly, we found that there was significant difference in the performance accuracies when relying on the data from one laboratory compared to the data from multiple laboratories (Figures 4A, B). With exception of a single worst performing classifier, SVM, the classifiers performed better on the RBM dataset, which came from a single study, than on the NCBI dataset, which was obtained by merging multiple independent studies. Moreover, this difference held for both the gene and isoform based models. Next, we evaluated if the prediction accuracy depended on the transcript counting approach. To do so, TCGA expression values were calculated based on (i) raw counts and (ii) log_2_ normalized counts with respect to the gene length and sequencing depth. The results showed that there was a strong preference, in terms of accuracy, in raw counts for the gene-based classifiers, but to a lesser extent for the isoform-based models. However, the opposite was observed where isoform-based models were more accurate when using log_2_ normalized counts (Figures 4C, D). There was less variability (less than 0.3 in the maximum difference of f-measure values across all methods for each classification task) when considering isoform-based versus gene-based models.

**Figure 4.**
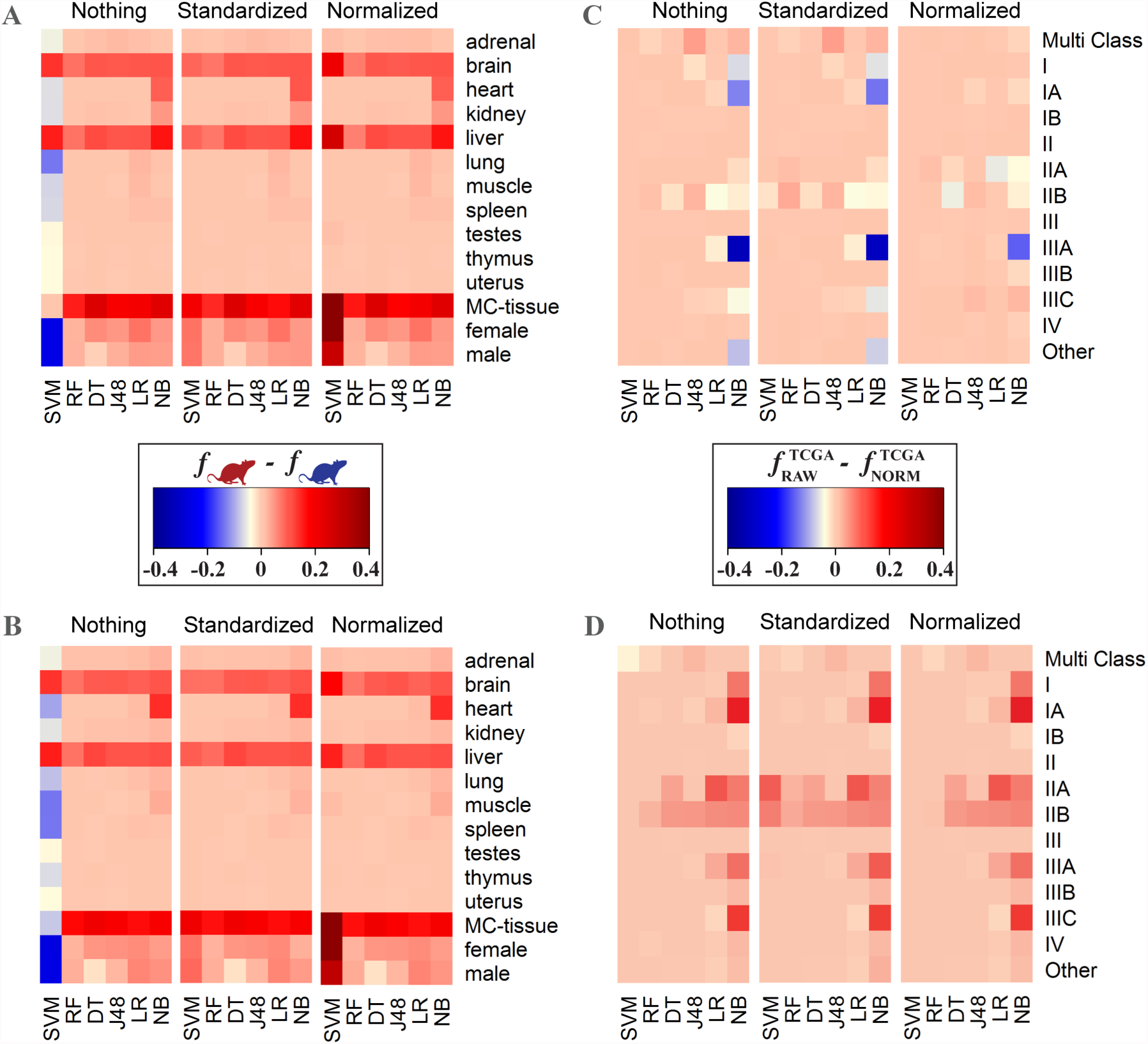
Heat map representation showing the influence of different factors on the accuracy performance. Panel (A) and (B) represent the difference in performance accuracies, calculated by f-measure, between RBM (single-lab) and NCBI (multi-lab) datasets for gene-based (A) and isoform-based (B) classifications, respectively. Panel (C) and (D) represent the difference in f-measure between the classifiers trained on the TCGA expression values quantified as either raw counts or log_2_ normalized counts with respect to gene length and sequencing depth, Shown are f-measure differences for gene-based (C) and isoform-based (D) classifications, respectively. The bottom x-axis represents the machine learning techniques (DT = Decision Table, J48 = J48 Decision Tree, LR = Linear Regression, NB = Naïve Bayes, RF = Random Forest, SVM = Support Vector Machine). The y-axis represents the classes where MC stands for multiclass. The top x-axis represents normalization techniques including Nothing (no normalization), Standardized, and Normalized.

**Figure 5.**
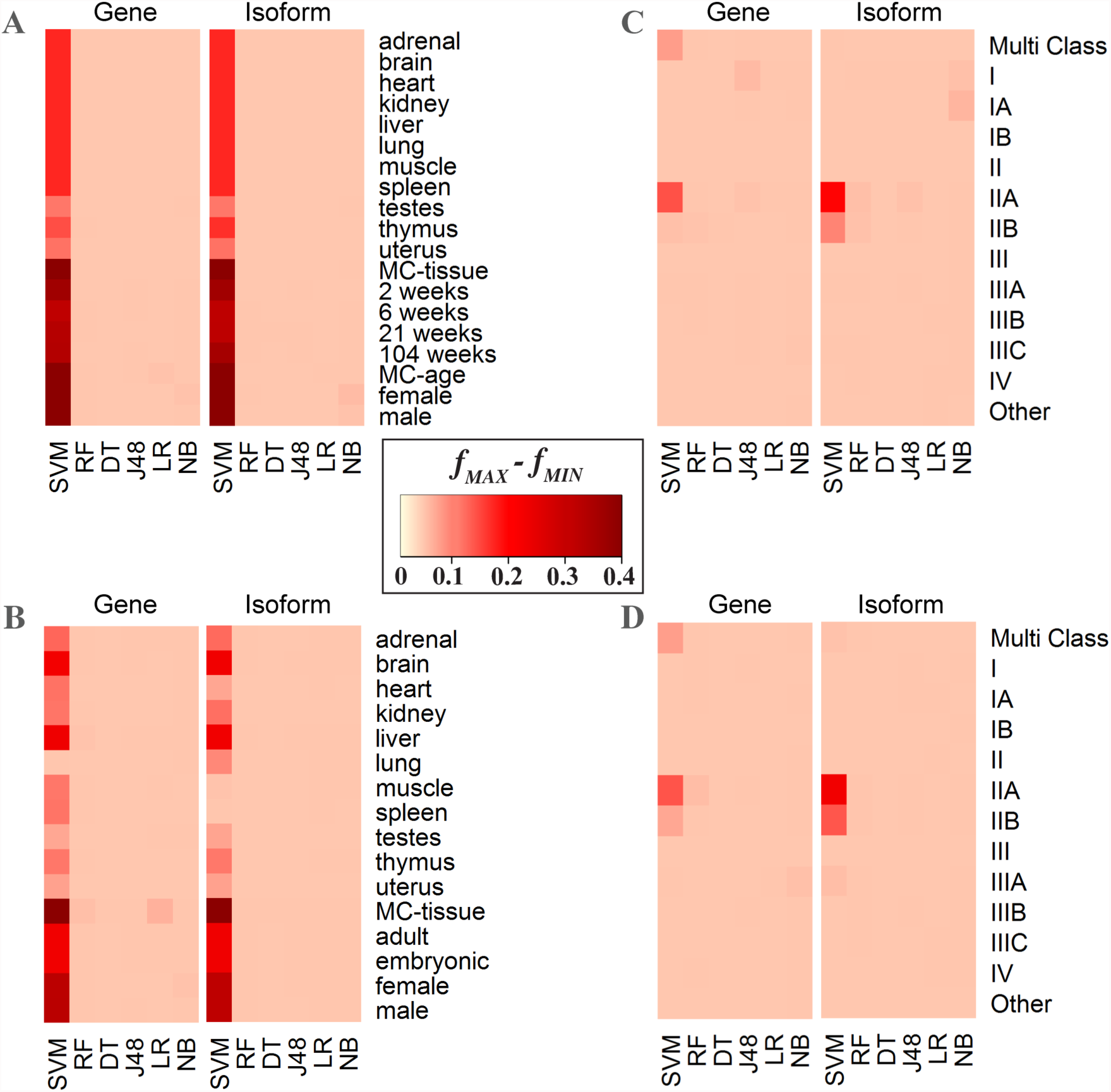
Heat map representation showing the influence of normalization techniques on the performance accuracy. To demonstrate the variability attributed to machine learning normalization technique, the intensity of the color represents the difference between the maximum and minimum *f*-measures achieved for a specific classification task and specific classifier across all three normalization protocols. The x-axis represents the machine learning techniques (DT = Decision Table, J48 = J48 Decision Tree, LR = Linear Regression, NB = Naïve Bayes, RF = Random Forest, SVM = Support Vector Machine). The y-axis represents the classes where MC stands for multiclass. The upper x-axis stands for whether the difference is from gene or isoform expression value. SVM is the only method that has significant changes due to normalization technique.

Finally, we considered different normalization techniques across the gene- and isoform-based classifiers. The general trend observed was little to no difference in performance accuracies when either different normalization protocols or no normalization at all were used. The main difference was observed on the IIA and IIB classification tasks with the classifiers trained on the TCGA dataset. Interestingly, irrespective of the gene- or isoform-based data, the SVM classifiers had the highest variability in classification accuracy (Fig 5).

### Normal Phenotype Classification Tasks: Age, Sex, and Tissue Classification of Rat Samples

The Rat Body Map (RBM) represents a dataset with the least amount of noise due to non-biological variation becomes it comes from a single laboratory, which uses the same sample and library preparation protocols and a fixed sequencing depth. From this dataset we identified eleven tissue types, four age groups, and both sexes. We then defined 17 “one-against-all” binary classification problems (Figure 3). Additionally, we merged the tissue and age groups and applied a multiclass classifier. As it was previously mentioned, the normalization technique had little effect on the classification results except for SVM where the classification accuracy difference was as high as 50% (Figure 5).

For the tissue classification, including multiclass tissue classification, the models achieved 100% accuracy and 1.0 f-measure based on the assessment protocol and irrespective of the machine learning method. However, when considering normalization technique, SVM had the accuracy ranged between 75.3% to 99.8% and 0.39 to 1.00 f-measure. The age group classification represented a more difficult task, with the classification accuracy ranging between 40.2% to 100% and f-measure ranging from 0.40 to 1.00. For the 2-week and 104-week age groups, the classifiers again achieved nearly 100% accuracy and 1.0 f-measure across all machine learning techniques. The 6-week and 21-week age groups were predicted with over 97% accuracy using random forest, j48, and logistic regression classifiers, while naïve Bayes could only achieve 81.1% and SVM with 40.2%. Similar pattern was observed in sex classification, where logistic regression and random forest achieved more than 97.3% in accuracy, but naïve Bayes could reach only 86.1%.

The NCBI dataset was expected to result in a greater variation of the feature values, compared with the RBM dataset, since it included the data from multiple research laboratories that sequenced different rat samples and even strains using different library preparation protocols. The same types of classification tasks were considered, including tissue, age, and sex. Since this dataset represents all publically available data in rat obtained using the same sequencer model, it included more tissue types than the RBM dataset. For consistent comparison, only those tissue types that were previously included in the RBM dataset were chosen for the NCBI dataset for the binary classification. However, for the multiclass tissue classification problem, the labels were determined based on the entire range of organs and tissues that the samples originated from, thus including more tissue types than in the RBM dataset. In contrast, the age group classification for the NCBI dataset was more limited than that one for the RBM dataset, since some samples in the former did not include the detailed age information. Therefore, the age types for the NCBI datasets were reduced to either adult or embryonic types.

The RNA-Seq data normalization did not have an effect on the classification results for the NCBI dataset: the performance difference when using the normalized and unnormalized datasets was only observed for the SVM classifier, the method that performed the worst out of the six supervised learning methods. The binary tissue-based classification performed well overall, reaching over 99.7% in accuracy and 0.99 in f-measure for the top-performing random forest classifier. Interestingly, the worst performing classifier, SVM, achieved the accuracy of only 21.2% and 0.07 f-measure for the gene-based tissue multi class. The analysis of method performances for multiclass classification tasks revealed that classification of several tissue types was particularly challenging for some of the less accurate methods. The binary tissue type classification tasks reporting the lowest accuracies included brain and liver tissue classification, with 79.1%-94.3% in accuracies and 0.70-0.94 in f-measure values, depending on the supervised learning method used. For the harder problem of multiclass tissue classification, the performance of the classifiers was highly variable, with the accuracy ranging from 0.7% to 84.3% and f-measure from 0.07 to 0.84, and with the observation that the random forest classifier was, again, the best performing method. Differentiating between embryonic and adult samples as well as between the sexes were easier tasks compared to the tissue origin. The age classification accuracy ranged from 83.2% to 98.3% and f-measure from 0.75 to 0.97 across all six supervised learning. The sex classification task had classification accuracy ranging between 71.1% and 97.3% and f-measure between 0.59 and 0.97. Interestingly, the consistently poor performance of the SVM classifier was not dependent on the normalization technique.

### Disease Phenotype Classification Tasks: Breast Cancer vs, Healthy and Stage Classification of Human Samples

Based on the promising results for the normal phenotype classification tasks, we further increased the difficulty of classification task by predicting different pathological stages of breast cancer using gene-based and isoform-based data. To evaluate if this classification task could benefit from additional information, we assessed the method performances based on the RNA-Seq data with log_2_ normalization in addition to the three types of normalization used in the two previous classification problem. The classification performance was heavily dependent on the supervised learning method with accuracies ranging from 20.2% to 99.8% and f-measure ranging from 0.21 to 1.00, and with naïve Bayes and SVM classifier being the worst performing classifiers. Furthermore, when considering all classes and log_2_ normalization, the accuracies decreased by as much as 60%, and the only method that benefited from the normalization was the poorly performing SVM classifier.

For each stage of breast cancer, we were able to achieve at least 78.3% in accuracy and 0.77 in f-measure. However, there is a significant variability within all parameters tested (Figure 2C & D). Similar to the analysis for the RBM and NCBI datasets, random forest had the highest performance across all stages of breast cancer based on 71.3% to 99.8% accuracy and 0.64 to 1.00 f-measure. The most difficult stages to classify were stages IIA and IIB, with the average difference in accuracy between 21.3% accuracy and 0.29 f-measure. Unlike the RBM and NCBI datasets, there were classes, such as Stage IIB, where naïve Bayes and SVM outperformed random forest by 5% in accuracy and 0.11 in f-measure. The easiest stages to classify were stages II and III with 99.8% accuracy and 1.00 f-measure.

In contrast to the RBM and NCBI datasets, the worst performing models for each class were highly variable, depending on the parameters chosen. For example, for stage IIA, the logistic regression classifier was the best performing model at 78.2% accuracy and 0.77 f-measure. However, the worst performing model was J48 at 60.1% accuracy and 0.60 f-measure. Similarly, for stage I the worst performing classifier was naïve Bayes with 53.7% accuracy and 0.63 f-measure, while the best performing classifier was random forest with 91.4% accuracy and 0.84 f-measure. On the other hand, for stage III binary classification the performance was 99.5-99.8% accuracy and 0.996-0.998 f-measure across all classifiers and parameter sets. These results demonstrated that no single method and parameter set was able to always outperform all others.

## Discussion

This work achieves two aims. The first aim is to broadly assess how well the supervised machine learning methods perform in various biological classifications by utilizing exclusively the RNA-Seq data. This aim is supported by our rationale that the key biological patterns should be recoverable from the transcriptomics data. Our second aim is to investigate whether relying on the isoform-level expression, which provides details on the alternatively spliced isoforms, can increase the accuracy of biological classification compared to the gene-level expression. Since the data patterns detected by the machine learning techniques during their training stage are highly dependent on the type of biological classification, we wanted for our assessment to cover multiple aspects. Specifically, we evaluated the performance of six widely used supervised classifiers across different RNA-Seq datasets, organisms, and normalization protocols, totaling in 61 classification problems and 2,196 individual classification tasks. Each task separately utilized the gene-level and isoform-level expression datasets. The main purpose behind our study was to demonstrate the importance of enriching RNA-Seq data with the differentially expressed transcripts for the biological classification tasks, suggesting that limiting the RNA-Seq analysis to the differentially expressed genes would, in turn, limit the capabilities of machine learning algorithms. As a result, several important conclusions were made.

First, we found that the accuracy of machine learning classifiers depended on how much data variation associated with the type of sequencers, library preparation, or sample preparation was introduced. Our rat datasets were specifically selected to compare the differences in data variation and in classification accuracies. The first dataset (RBM) was chosen because it included samples representing multiple age groups, tissue types, as well as sex [36], while these data were generated by only one research group and using the same sequencer. Thus, possible variation due to the type of sequencers or preparation protocols was expected to be minimal. Furthermore, we downloaded and processed the raw RNA-Seq reads using our in-house protocol and thus excluding possible variation due to different RNA-Seq analysis techniques. Our second dataset (NCBI) incorporated all publically available RNA-Seq data for rat using the same sequencer model, thus minimizing possible sequencer-based bias, a well-documented source of variation [42]. The NCBI dataset included 29 studies from multiple laboratories and represented the same classes as in the RBM except for the age groups. As expected, higher variation negatively affected the accuracy across predominantly all machine learning methods, normalization protocols and classification tasks. On the other hand, even for the NCBI dataset, the accuracies for all top-performing binary classifiers were never below 90% either for gene-level or for isoform-level expression data, suggesting minimal influence of the batch effect on the supervised classifiers.

Second, our study suggested that the standard data normalization techniques were not needed for RNA-Seq data except when using the poor-performing SVM classifiers. Random forest and logistic regression classifiers performed consistently well with each of the normalization technique but also without them, regardless of the classification task. However, there were several normalization techniques that were specific to RNA-Seq data, including RPKM (reads per kilobase per million reads), FPKM (fragments per kilobase per million reads), and TPM (transcripts per kilobase per million reads) [3]. Assessing whether these normalization techniques have an effect on classification accuracy should be considered for future studies.

Third, we found that the overall performance of the most accurate machine learning classifiers was very strong, with a few exceptions. In fact, for several classification tasks including all tissue classes, 2 week, and 104 week from RBM dataset, stage I from TCGA dataset, and the top-performing classifiers achieved a perfect 1.0 f-score, while for the majority of other tasks, the accuracy and f-measure were no less than 0.9 and often achieved by more than one classifier. From the biological perspective, it was surprising to see how well the classifiers performed on the normal phenotype datasets, in spite of significant variations in the sample and library preparation by different labs as well as the difference in rat strains. Intuitively, the expression values should have high variability due to these differences. The few exceptions in excellent performance were the multiclass age group classification for the normal phenotype datasets and classifications of clinical stages I, IIA, IIB and IIIA for the disease phenotype dataset, with stages IIA and IIB performing significantly worse. The clinical definition of IIA and IIB are based on the size of the tumor as well as evidence of cancer movement, and the reduced performance on each of these stages suggests that while there is a phenotype difference there may not be a strong molecular expression difference, which would cause a higher error rate by a classifier. The results also suggested that, from the diagnostic perspective, a more accurate AJCC classification methodology to distinguish those two phenotypes might be required to improve the stage prediction accuracy. The most consistent in the overall performance across all tasks were the random forest classifiers, which had been previously shown to perform exceptionally well for a number of bioinformatics tasks [43] and can be suggested as a reliable first choice for a biological classification task. Overall, our findings provided strong evidence that the supervised learning approach is readily available for the majority of the biological classification tasks.

Finally, we found that the classifiers that leveraged the isoform-level expression never performed worse and often outperformed the classifiers that used the gene-level expression data. For the normal phenotype tasks, the most profound difference was when considering the most challenging classification task—the multiclass classification of age groups. For the disease phenotype tasks, the most significant difference in performances of the classifiers that used gene-based and isoform-based expression data was again for the most challenging classes, the clinical stages IIA and II B of breast cancer. The better performance for the classifiers on the isoform-level data seems to be the expected result because the methods are trained on the enriched data, from the biological point of view. However, we note that the isoform-level data provides a significantly higher number of initial features, which could result in adding more noise to or potential overfitting of a classifier. Hence the importance of the feature selection and thorough model evaluation, which in this work suggests that the isoform-level information is a better choice when developing a biological classifier. Given that the isoform extraction methods continue to improve [44, 45], we expect further improvement in the accuracy of isoform-level based classifiers.

In conclusion, this study demonstrates that a supervised learning method leveraging isoform-level RNA-Seq data is a reliable approach for many biological classification tasks. It is a fast computational approach and can be fully automated for the projects that involve massive volumes of sequencing data and/or high number of samples. With the rapid advancements of RNA sequencing technologies as well as with continuous improvement of the isoform prediction methods, the accuracy of the machine learning approaches will be only increasing. We also expect for these methods to tackle more challenging tasks such as cell type classification, disease phenotype classification of common and rare complex diseases, and clinical stage classification across all major cancer types. Finally, we expect for advanced machine learning approaches, such as semi-supervised learning [46], deep learning [47], and learning under privileged information [48] to step in.

## Materials & Methods

The methodology used in this study compares three RNA-Seq datasets, six supervised machine learning methods, three normalization techniques, two RNA-Seq analysis pipelines, and 61 classification problems in order to assess if the features derived from the expression data at the alternative splicing level (*i.e*., isoform-based) can result in a higher classification accuracy than the features derived from the gene-based expression levels. Our approach attempts to systematically evaluate the classifiers that relied on these features from multiple perspectives, with a goal to provide a comprehensive analysis. We use the increasingly difficult biological classification tasks to assess the performances of classifiers in the presence of noise due to the difference in the biological sources, sequencers, and preparation protocols. The analysis is based on three RNA-Seq datasets, two from rat and one from human. The six supervised machine learning methods tested in this work include support vector machines (SVM), random forest (RF), decision table, J48 decision tree, logistic regression, and naïve Bayes. The three normalization protocols used include (1) pipeline-specific RNA-Seq count with no post-normalization, (2) pipeline-specific RNA-Seq count with normalization from 0 to 1, and (3) pipeline-specific RNA-Seq count protocol with standardization with respect to standard deviation. The two RNA-Seq analysis pipelines in this work, each employing different RNA-Seq count methods were the standard Tuxedo suite and RSEM. The 61 classification problems include binary and multiclass classifications of tissue types, age groups, sex, as well as clinical stages of breast cancer.

### Data Sources

Three datasets are used to demonstrate the usability of the isoform-level expression data for the supervised classification. The first two datasets are from rat samples of normal phenotype; the raw RNA-Seq data for both datasets is processed using our in-house protocol. The last dataset consists of already processed RNA-Seq data from human breast cancer samples [49]. The first, RBM, dataset is obtained from the Rat Body Map and includes 660 samples from 12 different rats from the F344 rat strain [36] covering 4 different age groups, 11 tissues, and both male and female rats. Publicly available raw mRNA RNA-Seq data from the Rat Body Map (http://pgx.fudan.edu.cn/ratbodymap/) is downloaded and processed for the gene and isoform levels of expression. The second dataset, NCBI, includes all publically available raw RNA-Seq data from rat samples that are sequenced using Illumina Hi-Seq 2000 and available from the NCBI GEO DataSets collection (http://www.ncbi.nlm.nih.gov/gds, Suppl. Table S3). In total 1,308 samples are used, which represents 29 different projects. In contrast to the processing of the data for the first dataset, these 29 projects used a variety of library preparation protocols and adapters to process their samples. The third, TCGA, dataset is obtained from The Cancer Genome Atlas data repository [49] and includes 1,216 breast cancer patients diagnosed with different pathological cancer stages (as defined by the American Joint Committee on Cancer, AJCC [39]). The class distributions for all datasets are shown in Suppl. Figures S9-S11.

### RNA-Seq Pipeline

RNA-Seq analysis encompasses three main stages: preprocessing, alignment, and quantification. There are a number of methods to perform each of these three basic steps, while the debate on the most appropriate methodology continues [3]. In this work, we expect for the variation due to data processing to be minimal since the same processing pipeline is used for each dataset. Two different RNA-Seq pipelines are implemented and applied to each dataset for both gene and isoform levels of expression. These two pipelines leverage different algorithms and different metrics [50]. For the RBM and NCBI dataset, all raw RNA-Seq data are downloaded from the SRA repository (https://www.ncbi.nlm.nih.gov/sra) using unique project IDs (Suppl. Table S3). SRA file formats are then converted into fastq format. These files are used as input for the preprocessing stage. The preprocessing is done using Fastx Tools with the settings that removed reads shorter than 20 bp. All nucleotides with quality scores of less than 20 are converted into N’s [51]. The alignment is done against the rat genome version rn5 [52] using Tophat v2 and its default settings [53]. Quantification for both gene- and isoform-based expression levels is performed using Cufflinks v2 [54] and Ensembl transcript annotation v75 [55]. The Cufflinks is set to use the transcript annotation for quantification with other settings being default. For the TCGA dataset, MapSlice [56] is used for alignment and RSEM [57] for quantification. The final output includes expression levels for each sample at both gene and isoform levels. We note that the gene-based expression values are the summation of all isoforms determined to be associated with the corresponding gene.

### Supervised Learning Classifiers

The quantified expression values obtained from Cufflinks are then used to train and assess six supervised classifiers for each task. Two types of classification tasks are considered: one-against-all and multiclass. Our classification approach leverages feature-based supervised learning methods. Each post-processed RNA-Seq sample is represented as a feature vector, where each feature represents the isoform- or gene-level expression value for a specific gene or AS isoform corresponding to this gene. Expression samples may vary in length, thus to generate feature vectors of the same length, we compute the intersection of all samples in terms of the feature set that represents each sample. We next rank the importance of each feature and select subsets of the features that best describe their respective classes using the Best First (BF) feature selection method [40]. The BF method is driven by the property that the subsets of important features are highly correlated with a specific class and are not correlated with each other. The method is described as a greedy hill climbing algorithm augmented with a backtracking step, where the importance of features is estimated through one-by-one feature removal. All machine learning methods are implemented using the Weka package version 3.7.13 [58].

Due to a large number of features for genes and even greater number of features for the isoforms (~20,000-73,000) using the base classification and even BF method was not computationally feasible, thus the modification to the original methodology is implemented allowing to reduce the processing time. The modifications includes introducing multiple splits of the features followed by two rounds of BS feature selection. Specifically, we split the data into 1,000 subsets and perform feature selection on each subset independently. After feature selection is performed on all splits, the selected features are merged, and another round of feature selection is performed. Our solution reduces the time needed to compute from several weeks to hours and still able to successfully select a reduced feature set that allows for accurate classification.

### Machine Learning Technique Rationale

A broad selection of supervised learning approaches were implemented to test whether performance could be improved, depending on the method tested. The machine learning methods have different assumptions on how the data are structured; the methods also vary in their treatment of the class outliers and convergence w.r.t number of training examples.

The first two classifiers, naïve Bayes and logistic regression, are often regarded as the baseline methods due to their simplicity and robustness. *Naïve Bayes* classifier is a probabilistic method that has been used in many applications including bioinformatics and text mining [59-63]. It is a simple model that leverages the Bayes rule and describes a class of Bayesian networks with assumed conditional independence between the numerical features. The use of this “naïve” assumption makes the method computationally efficient during both the training and classification stages. Furthermore, while the probability estimation by naïve Bayes is reported to be not very accurate [64], a threshold-based classification performance is typically very robust. In our implementation, the numeric estimator precision values are chosen based on analysis of the training data and is set to 0.1 The batchSize parameter that specifies the preferred number of instances to process during training if batch prediction is being performed is set to 100. *Logistic regression* is another type of a simple machine learning classifier that has been compared with naïve Bayes in terms of accuracy and performance [65]. Different versions of logistic regression models are often used in bioinformatics applications [66-69]. In this work, we implemented a boosting linear logistic regression method without regularization and with the optimal number of boosting iterations based on cross validation.

The next three classifiers, decision tables, J48, and random forest, are the decision tree based algorithms. A *decision table* is a rule-based classifier commonly used for the attribute visualization and less commonly for classification. The rules are represented in a tabular format using only an optimal subset of features that are included into the table during training. The decision table is a less popular approach for bioinformatics and genomics classification tasks, however it has showed a superior performance in some bioinformatics applications [70], and therefore is included into the pool of methods. The decision table model is implemented as a simple majority classifier using the Best-First method for searching. *J48* is an open source implementation of perhaps the most well-known decision tree algorithm, C4.5 [71], which is, in turn, is an extension of Iterative Dichotomiser 3 (ID3) algorithm [72]. C4.5 uses the information-theoretic principles to build decision trees from the training data. Specifically, it leverages the information gain and gain ratio for a heuristic splitting criterion with a greedy search that maximizes the criterion. Furthermore, the algorithm includes a tree pruning step to reduce the size of the tree and avoid the overfitting. In this work, the implementation of J48 was done with the default confidence threshold of 0.25 and minimum number of instances per leaf set to 2. *Random forest* is an ensemble learning approach, where many decision trees are generated during the training stage, with each tree based on a different subset of features and trained on a different part of the same training set [73]. During the classification of unseen examples, the predictions of the individually trained trees are then agglomerated using the majority vote. This bootstrapping procedure is found to efficiently reduce the high variance that an individual decision tree is likely to suffer from. The random forest methods have been widely used in bioinformatics and genomics applications due to their versatility and high accuracy [73]. In this work, due to a large but highly variable number of features the number of attributes, *K*, randomly selected for each tree is dependent on the classification task and is defined as *K* = [log_2_ *n* + 1], where *n* is the total number of features. The number of sampled trees per each classifier is set to 100.

The last method, Support Vector Machines (SVM) represents yet another family of the supervised classifiers, the kernel methods [74]. It is among the most well-established and popular machine learning approaches in bioinformatics and genomics [26, 75-77]. SVM classifiers range from a simple linear, or maximum margin, classifier where one needs to find a decision boundary separating two classes and represented as a hyperplane, in case of a multi-dimensional feature space, to a more complex classifier represented by a non-linear decision boundary through introducing a non-linear kernel function. For our SVM model training, Radial Basis Function (RBF) was used, a commonly used kernel. The two parameters, *Gamma* and *C*, were set to 0.01 and 1, respectively.

### Training, Testing, and Assessment of classifiers

To evaluate each of the classifiers, a basic supervised learning assessment protocol is implemented. Specifically, the training/testing stages are assessed as a 10-fold stratified cross validation to eliminate the sampling bias. This protocol is implemented using Weka [58]. The reported result of assessment is based on the average *f*-measure for the 10-folds for testing dataset. *f*-measure incorporates recall (*Rec*, also called sensitivity) and precision (*Pre*) into one reported metric:

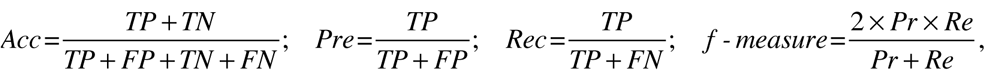

where *TP* is the number of true positives (correctly classified as class members for a specified class), *TN* is the number of true negatives (correctly classified as not class members), *FP* is the number of false positives (incorrectly classified as class members), and *FN* is the number of false negatives (incorrectly classified as not class members). While each of the above four measures are commonly used to evaluate the overall performance of a method, we primarily focus on the most balanced metric, *f*-measure, due to a high number of classification tasks to be reported.

## Availability

The supervised machine learning methods were implemented using the Weka platform (http://www.cs.waikato.ac.nz/ml/weka/). Data used are publically available from Rat Body Map (http://pgx.fudan.edu.cn/ratbodymap/), Geo Datasets (http://www.ncbi.nlm.nih.gov/gds), and the Cancer Genome Atlas (https://tcga-data.nci.nih.gov/tcga/tcgaHome2.jsp).

## Acknowledgements

The computations were performed on the HPC resources at the University of Missouri Bioinformatics Consortium (UMBC) and Worcester Polytechnic Institute (WPI). Additionally, would like to thank Dr. Lane Harrison for useful suggestions about the biological data visualization.

